# Interplay between MIG-6/papilin and TGF-β signaling promotes extracellular matrix remodeling and modulates the maintenance of neuronal architecture

**DOI:** 10.1101/2025.05.08.652809

**Authors:** Malika Nadour, Robert I. Valette R. L., Noémie Frébault, Valérie Fontaine, Lise Rivollet, Claire Y. Bénard

## Abstract

Neuronal architecture laid out during embryogenesis persists lifelong, ensuring normal nervous system function. However, the mechanisms underlying the long-term maintenance of neuronal organization remain largely unknown. We previously uncovered that the conserved extracellular matrix protein MIG-6/papilin impacts collagen IV remodeling and neuronal maintenance, such that disruption of MIG-6/papilin leads to a collagen IV fibrotic state and altered tissue biomechanics, thereby stabilizing neuronal architecture. Here, we combine incisive molecular genetics and *in vivo* quantitative imaging to determine how this *mig-6*-dependent fibrotic phenotype is modulated, by investigating the implication of the TGF-β pathway, which is well known to regulate fibrosis in mammals. Our findings highlight a mechanism whereby the interplay between MIG-6/papilin and the TGF-β pathway regulates ECM composition and neuronal maintenance, with MIG-6/papilin acting as a positive regulator of TGF-β signaling. This work provides key insights into the molecular basis of sustaining neuronal architecture and offers a foundation for understanding age-related neurodegenerative and fibrotic conditions.

## INTRODUCTION

Proper nervous system function depends not only on the accurate embryonic assembly of its architecture but also on its long-term preservation, as neuronal structures must withstand postnatal growth, maturation, and mechanical stresses from movement or injury [1–3]. Failure to maintain neuronal organization can compromise neuronal connectivity and function, and indeed, many neurodegenerative diseases are characterized by destabilized neuronal processes and synaptic loss [4–6]. Therefore, understanding how nervous system architecture is sustained throughout life is of critical importance.

Although crucial, the molecular mechanisms that protect neuronal architecture over time remain poorly characterized. Growing evidence points to the extracellular matrix (ECM) as a key player in maintaining tissue architecture in the nervous system. In the mature mammalian brain, the ECM constitutes up to 20% of total volume and is organized in both diffuse and condensed forms—such as basement membranes and perineuronal nets— each contributing distinct roles in synaptic plasticity, regenerative capacity, and neuronal stability [7–9]. Alterations in ECM composition and organization have been implicated in neurodevelopmental disorders, neurodegenerative conditions, age-related cognitive decline, and brain injury, underscoring the pivotal role of the extracellular environment in both physiological and pathological contexts [5, 10]. Still, our understanding of how ECM remodeling supports long-term neuronal architecture, particularly *in vivo* in the mature nervous system, remains limited.

More broadly, the ECM is subject to continuous remodeling, marked by dynamic changes in protein content, activity, and modifications such as crosslinking [11, 12]. Dysregulated ECM remodeling contributes to fibrosis, characterized by excessive ECM deposition, notably of collagens [13–15]. Transforming growth factor-β (TGF-β) is recognized as the most potent inflammatory mediator driving fibrosis [13]. The TGF-β superfamily comprises highly conserved signaling ligands, including vertebrate bone morphogenetic proteins (BMPs), *Drosophila* Dpp, and *C. elegans* DBL-1 (decapentaplegic/bone morphogenetic protein-like-1) [16–18]. The TGF-β signaling pathway, particularly the Sma/Mab pathway, is also evolutionarily conserved from invertebrates to humans [19]. This pathway initiates when BMP ligands bind to transmembrane serine–threonine kinase receptors, triggering phosphorylation of type I receptors by type II receptors. The intracellular signal is propagated as receptor-activated Smads (R-Smads) are phosphorylated by the type I receptor. Phosphorylated R-Smads then associate with common-mediator Smads (co-Smads) and additional transcription factors to regulate the expression of target genes [20, 21]. In mammals, BMPs regulate diverse cellular processes across development and adult tissue maintenance [22], including ECM homeostasis and basement membranes remodeling [23]. Moreover, the TGF-β signaling pathway may also contributes to neuronal development and function [24, 25] by regulating ECM production [26–28]. The ECM mediates cell adhesion and signaling via cell-ECM interactions, providing cells with growth factors including TGF-β [29–31]. The physical properties of the ECM also affect neural cell behavior [29]. However, the specific roles of TGF-β in brain ECM are not well understood, and whether TGF-β could participate in the lifelong maintenance of neuronal architecture has not been investigated.

The nematode *Caenorhabditis elegans* offers a genetically tractable *in vivo* model to study ECM roles in maintaining neuronal architecture over an organism’s lifetime, including the effects of the TGF-β signaling pathway on ECM organization. Despite undergoing a near 100-fold increase in body size from larva to adult [32], the nervous system of *C. elegans* retains its precise architecture established during embryogenesis [33, 34]. Neuronal cell bodies are organized into ganglia, and axonal processes follow well-defined tracts, all ensheathed by specialized ECM-rich basement membranes, composed of complex networks of highly cross-linked proteins, including non-fibrillar collagens (e.g. collagen type IV), laminins, fibulins, nidogen, and heparan sulfate proteoglycans [35]. The highly stereotyped nervous system and associated ECM enable investigation of ECM contributions to neuronal structures under controlled genetic conditions. Owing to its transparency, genetic tractability, and availability of cell-specific tools, *C. elegans* allows direct observation and manipulation of neurons and ECM components throughout the lifespan.

In *C. elegans*, the TGF-β Sma/Mab signaling pathway operates similarly to its vertebrate counterpart, with the secreted ligand DBL-1 binding to a receptor complex composed of the type II receptor DAF-4 and the type I receptor SMA-6. Ligand binding triggers DAF-4 to phosphorylate SMA-6, which subsequently phosphorylates SMAD proteins. Phosphorylated receptor-regulated SMADs (R-Smads) SMA-2 and SMA-3 associate with the co-Smad SMA-4, allowing their accumulation in the nucleus and transcriptional regulation of target genes [21, 36, 37]. Documented transcriptional targets of the DBL-1 pathway in *C. elegans* include genes encoding cuticle collagens and ECM-modifying enzymes [38–41]. The *C. elegans* TGF-β Sma/Mab pathway regulates body size, male-tail morphogenesis, innate immunity, reproductive aging, aversive olfactory learning, among other processes [18, 23, 42–49], and has been implicated in neuronal guidance [50]. However, whether TGF-β plays a role in the long-term maintenance of neuronal architecture has not been explored.

Prior *in vivo* genetic studies using *C. elegans* has provided important insights into the mechanisms supporting the integrity of neuronal assemblies. Among these are immunoglobulin superfamily proteins, including SAX-7 [51–56], homologous to the L1 family of cell adhesion molecules in mammals, the ectodomain of the FGF receptor EGL-15 [57] and ECM-associated proteins such as DIG-1 [58, 59] and ZIG family members ZIG-3 and ZIG-4 [60, 61]. Loss-of-function mutations in these molecules result in neuronal disorganization emerging late in development, well after normal morphology is established. Roles for the ECM and associated molecules in modulating synaptic maintenance have also been identified in *C. elegans*, with ZIG-10, collagen IV and metalloproteinases ADAMTS/GON-1 and MIG-17/ADAMTS sustaining synapse morphology [62–66].

Through a screen for suppressors of the neuronal disorganization defects in *sax-7* mutants, we previously identified the conserved ECM protein MIG-6/papilin [67], structurally related to ADAMTS proteases. We demonstrated that MIG-6/papilin cooperates with the ECM remodeling protease MIG-17/ADAMTS to regulate collagen IV, a major basement membrane component, highlighting the importance of ECM remodeling in maintaining neural stability. In *mig-6* mutants, collagen IV accumulates excessively and forms fibrotic structures, altering tissue biomechanics, and leading to stabilization of neuronal architecture [67]. Conservation of these molecules across species - for instance, L1CAM’s role in sustaining adult synapses and behavior in mice [68, 69] suggests shared principles underlying nervous system preservation from worms to mammals.

In this study, we investigated how the fibrotic-like ECM phenotype of *mig-6* mutants arises by examining the role of the TGF-β signaling pathway, a key regulator of ECM remodeling and fibrosis in vertebrates. We demonstrate that TGF-β signaling modulates the fibrotic ECM phenotype in *mig-6* mutants: inhibition of TGF-β exacerbates the collagen IV fibrotic phenotype and enhances the suppression of *sax-7* neuronal defects, while overexpression of TGF-β pathway components reduces this suppression. These findings suggest that a balance between MIG-6 function and TGF-β signaling fine-tunes collagen IV homeostasis and neuronal stability. Moreover, we show that other ECM components, including laminin and fibulin, contribute to this neuronal maintenance mechanism. Altogether, our work provides new insights into the interplay between ECM remodeling and a conserved signaling pathway, TGF-β signaling, in maintaining nervous system architecture and offers potential parallels to fibrotic mechanisms implicated in aging and neurodegenerative diseases.

## RESULTS

### The TGF-β pathway interacts with the ECM modulator *mig-6/papilin* in neuronal maintenance

We previously uncovered that a modified state of the ECM surrounding neuronal structures positively impacts the long-term maintenance of neuronal organization [67]. Indeed, loss of *mig-6S*/papilin, a conserved ECM protein that impacts collagen IV remodelling [35, 67] suppresses the neuronal disorganization displayed by mutants of the *sax-7* neuronal maintenance gene [67]. Chemosensory neurons ASH and ASI, which can be readily visualized with reporter P*sra-6::*DsRed2 (**Fig. 1B**)[56], acquire a stereotypical positioning during embryogenesis that is preserved throughout life and constitute a reliable model to study the maintenance of neuronal architecture lifelong. In wild-type animals, the soma of neurons ASH and ASI are located posterior to the nerve ring (where their axons project). In *sax-7* mutants, whereas these neurons initially develop normally during earlier development, they later progressively become displaced from the 4th larval stage onward, with neuronal soma getting displaced and the nerve ring shifting posteriorly, which leads to the soma aligning with or being anterior to the nerve ring (**Fig. 1B,C**) [51, 54, 56]. Loss of function of *mig-6/papilin per se* does not alter neuronal organization, quite the opposite, it stabilizes neuronal organization in *sax-7* mutants (**Fig. 1B,C**), and also safeguards ASH/ASI architecture in otherwise wild-type animals subjected to the stressful condition of constant swimming [67].

**Figure 1.**
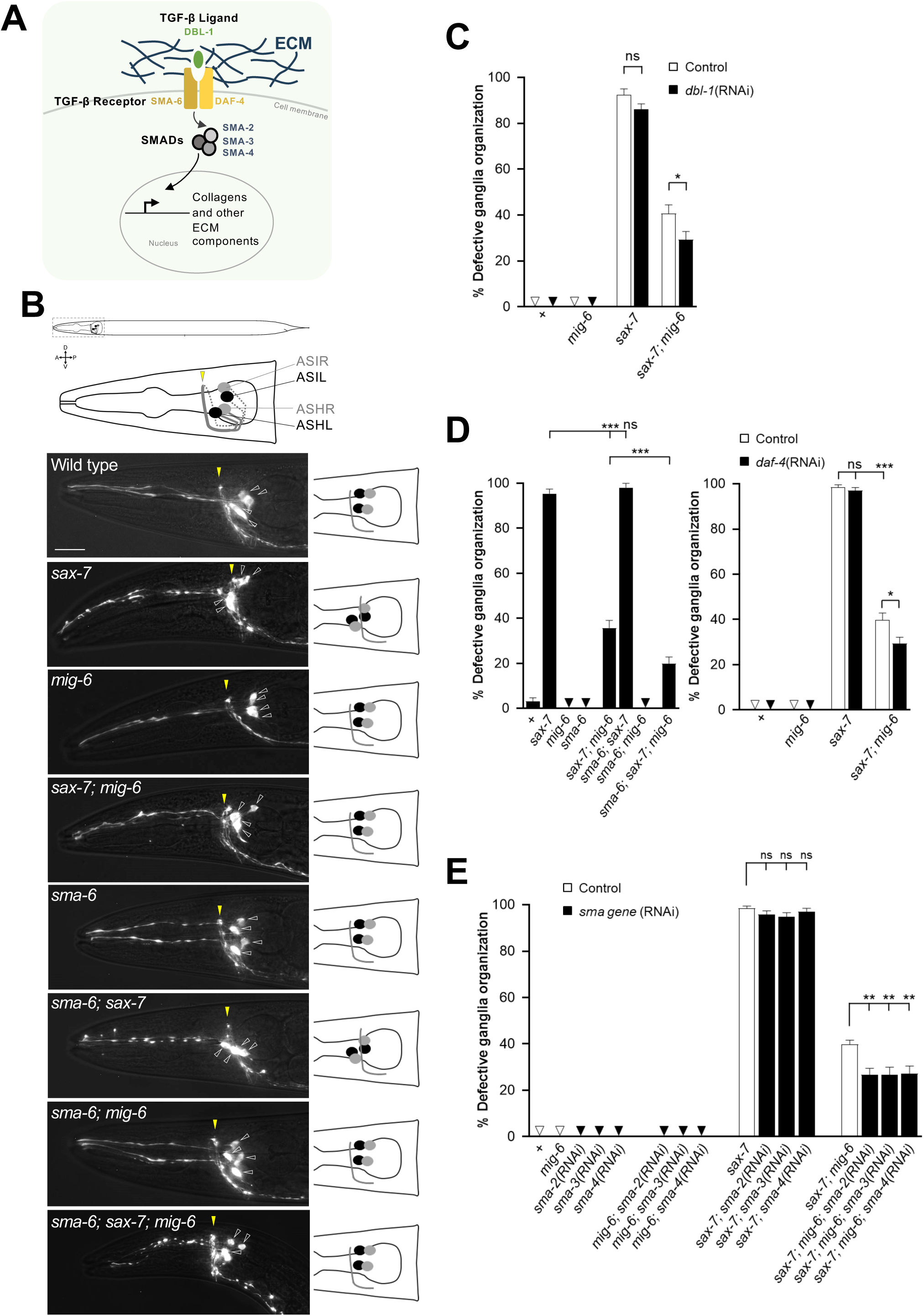
Disruption of the TGF-β pathway enhances the neuronal stabilization resulting from the loss of *mig-6/papilin*. **(A)** Schematics of the conserved TGF-β pathway (*C. elegans* protein names are indicated), which regulates ECM genes’ expression. **(B)** Fluorescence images of the head region of 2-day-old adults, where the chemosensory neurons ASH and ASI were visualized using fluorescent reporter P*sra-6::DsRed2* (schematized above and on the right). In the wild type, the four cell bodies (indicated by empty arrowheads; each ASH and ASI pair has a soma in the left ganglion and a soma in the right ganglion) are positioned posterior to the nerve ring (indicated by the yellow arrowhead) throughout life. In *sax-7* mutants, ASH and ASI soma, as well as the nerve ring, are initially positioned normally but by the 4th larval stage or older their relative positioning becomes progressively altered. Mutation *mig-6(qv33)* suppresses the neuronal disorganization of *sax-7* mutants. Scale bar, 20 µm. **(C)** Depletion of the TGF-β ligand DBL-1 by RNAi enhances the suppression of *sax-7* neuronal defects by *mig-6(qv33)*. Quantification of ASH and ASI displacement in 2-day adults of wild type, *sax-7(qv30)*, and *mig-6(qv33)* single and double mutant animals, treated with *dbl-1*(RNAi) [or control (empty RNAi vector)] since the L1 stage. **(D)** Loss of function of TGF-β type 1 receptor subunit SMA-6 and depletion of TGF-β type 2 receptor subunit DAF-4 by RNAi enhance the suppression of *sax-7* neuronal defects by *mig-6(qv33)*. Quantification of ASH and ASI displacement in 2-day adults of wild type, *sax-7(qv30)*, and *mig-6(qv33)* single and double mutants combined with the *sma-6(wk7)* mutation, or with *daf-4*(RNAi) treatment [or control RNAi (empty vector)] since the L1 stage. **(E)** Depletion of the TGF-β pathway downstream effectors SMADs by RNAi enhances the suppression of *sax-7* neuronal defects by *mig-6(qv33)*. Quantification of neurons ASH and ASI displacement in 2-day-old adults subjected to *sma-2*(RNAi), *sma-3*(RNAi), or *sma-4*(RNAi) [or control RNAi (empty vector)] since the L1 stage. Error bars are the standard error of the proportion. Comparisons were made with z-tests. **<u>Note</u>**: Throughout all figures of this work, asterisks denote significant difference (\**P* ≤ 0.05, \*\**P* ≤ 0.01, \*\*\**P* ≤ 0.001); n.s., not significant; appropriate *post hoc* tests were performed for multiple comparisons (see **Materials and Methods**); “+” indicates wild-type strain; the small triangles indicate 0% penetrance; all sample sizes and source data are provided in **Supplementary Information**.

The *mig-6/papilin* mutation stabilizing effect on neuronal architecture results from an altered ECM state, characterized by the build-up of major extracellular matrix component collagen IV, which accumulates as fibrotic forms [67]. The TGF-β/BMP pathway has been extensively implicated as a key modulator of fibrosis in vertebrates. We therefore asked whether the TGF-β pathway in *C. elegans* may also impact the state of the ECM particularly in relation with the maintenance of neuronal architecture. We focused on the *C. elegans* DBL-1/SMA-6/DAF-4 pathway as it has been documented to regulate mainly the collagen composition of a specialized ECM (of the cuticle encasing the animal’s the body), and because ECM collagens can, in turn, influence this signaling cascade [38, 70]. This TGF-β pathway is composed of evolutionarily conserved core components: the ligand (DBL-1/BMP), type I and type II receptors (SMA-6/RI and DAF-4/RII), and the R-Smads downstream effectors (SMA-2 and SMA-3), and the co-Smad (SMA-4) (**Fig. 1A**; [18, 48, 49, 71–74].

We explored whether this TGF-β pathway impacts neuronal maintenance by testing genetic interactions with *sax-7* and *mig-6S/papilin*. We knocked down components of the DBL-1/TGF-β pathway using RNA interference (RNAi) to deplete transcripts of *dbl-1*, *daf-4*, *sma-2*, *sma-3*, or *sma-4*, as well as using a null mutation in the gene *sma-6*, *wk7*. These genetic manipulations of the TGF-β pathway were combined with the null mutation *sax-7*(*qv30)* [56] and the well-characterized missense loss-of-function mutation *mig-6(qv33)* [67]; null *mig-6* mutants are sterile [75]. We found that inactivating the TGF-β pathway by *dbl-1*(RNAi) ligand knockdown, *sma-6(wk7)* receptor mutation, *daf-4*(RNAi) receptor knockdown, or knockdown of the downstream effectors by *sma-2(RNAi)*, *sma-3(RNAi)* or *sma-4(RNAi)* -in an otherwise wild-type background-did not impact the architecture of the ASH/ASI neurons (**Fig. 1B-E**). Thus, inactivation (by *sma-6*/receptor mutation) or downregulation (by RNAi) of the TGF-β pathway alone does not affect the development or position maintenance of these neurons. Similarly, the ASH/ASI neurons in animals lacking both TGF-β signaling and *mig-6*/*papilin* were also like the wild type (**Fig. 1B-E**). Further, knockdown of the TGF-β pathway in the *sax-7(qv30)* mutant background resulted in animals that are identical to *sax-7* single mutants (**Fig. 1B-E**), indicating that the TGF-β pathway does not modify the mutant neuronal phenotype associated with *sax-7*. In contrast, combining the inactivation of the TGF-β pathway with the *mig-6(qv33)* mutation resulted in an enhanced suppression of the *sax-7* neuronal maintenance defects compared to the suppression obtained by the *mig-6* mutation alone. Indeed, whereas in double mutant animals *sax-7; mig-6* the loss of function of *mig-6/papilin* suppresses *sax-7* defects from 90% to 40%, the combined disruption of the TGF-β pathway consistently leads to a more profound reduction of the neuronal defects down to 20-30% in triple loss of function animals *sma-6*; *sax-7; mig-6*, or in double mutant animals *sax-7; mig-6* treated with RNAi targeting any of the five TGF-β pathway components (**Fig. 1B-E**). These results reveal an impact of the TGF-β pathway on neuronal maintenance in a *mig-6*-dependent manner, with its loss amplifying or potentiating the stabilizing effect of the loss of function of *mig-6/papilin* on neuronal architecture. This raises the possibility that the TGF-β signaling pathway might be implicated in the mechanism through which *mig-6*/papilin modulates neuronal maintenance.

### *mig-6*-dependent collagen IV remodeling is affected by the TGF-β pathway

*mig-6*/papilin plays a key role in collagen IV homeostasis [35, 67], and we previously demonstrated that both levels and the reticulation of collagen IV in the head region -in the vicinity of neuronal structures under study-are key for neuronal maintenance [67]. Since TGF-β pathway disruption intensified the *mig-6-*mutation stabilizing effect on neuronal architecture (**Fig. 1**), we asked whether the altered collagen IV state of *mig-6* mutants might also further be exacerbated by the loss of TGF-β pathway function. To test this, we constructed mutant strains of *mig-6(qv33)* and *sma-6(wk7)* carrying reporter *qyIs46* P*emb-9*::EMB-9::mCherry [76] to visualize collagen IV in the head region. Compared to wild-type animals, *mig-6/papilin* mutants accumulate collagen IV at higher levels (**Fig.2 A,D**) [67], including in form of extracellular fibrotic-like structures (**Fig. 2A-C**) [67]. In contrast, we found that TGF-β receptor *sma-6* single mutants display collagen IV pattern and levels similar to the wild type (**Fig. 2A-D**). We next examined collagen IV in double mutants *sma-6; mig-6*, where analysis of the percentage of animals displaying fibrotic-like structures did not reveal differences with the already fully penetrant *mig-6* single mutants (**Fig. 2B**). We therefore quantified the number of fibrotic-like collagen IV structures per animal and found that the fibrotic phenotype significantly increases in *sma-6; mig-6* double mutants, compared to *mig-6* single mutants (**Fig. 2C**). Collagen IV fluorescence intensity also increases in *sma-6; mig-6*, compared to *mig-6* single mutants (**Fig. 2D**), indicating that while loss of TGF-β pathway signaling alone does not alter collagen IV, it heightens the already compromised state of collagen IV of *mig-6/papilin* mutants.

**Figure 2.**
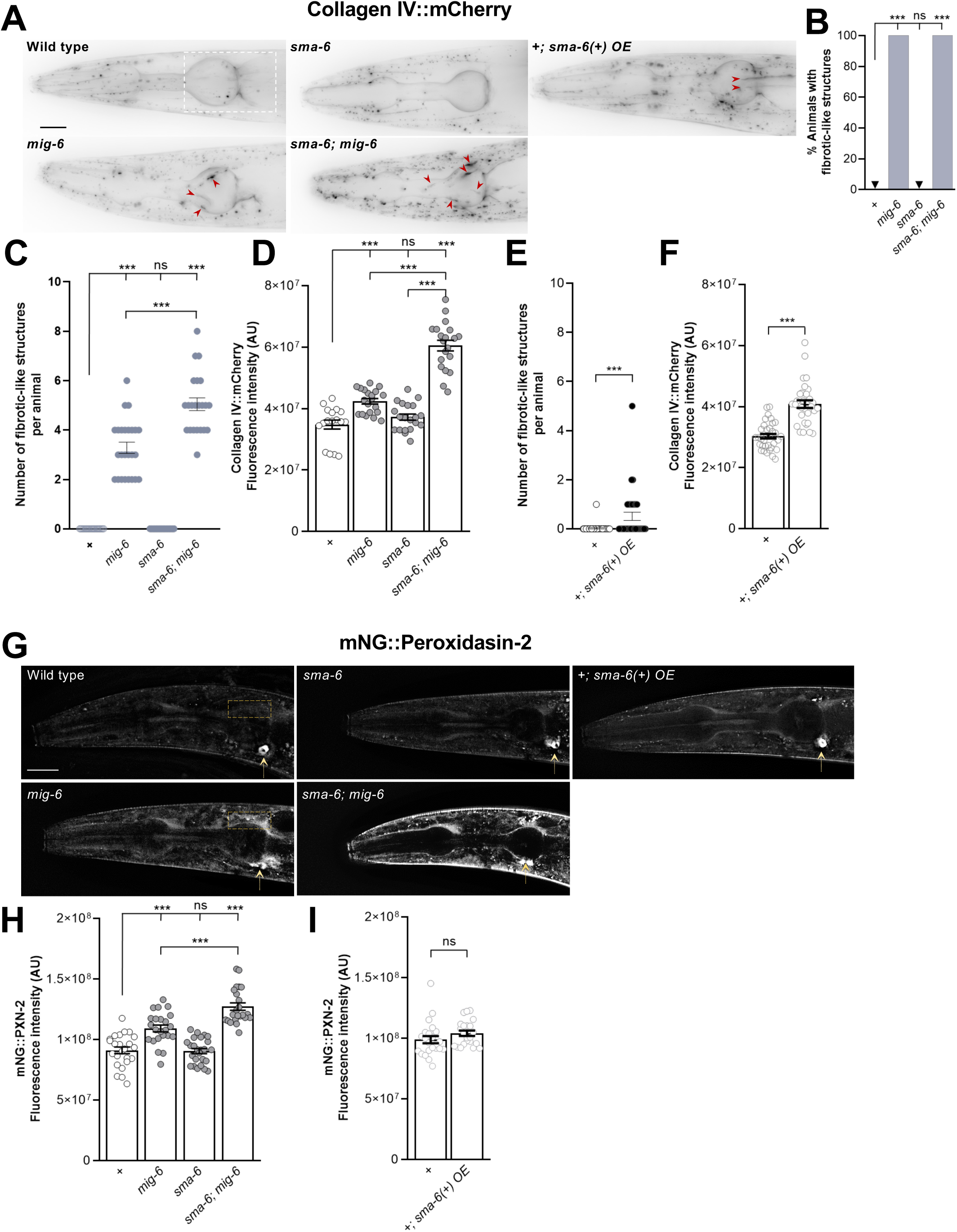
Altering TGF-β pathway function enhances the collagen IV fibrotic state of *mig-6/papilin* mutants. **(A-F)** Analysis of 2-day-old adult animals; collagen IV was visualized using *qyIs46* P*emb-9*::EMB-9::mCherry. **(A)** Fluorescence images (deconvoluted sum projections) of the head region, where ganglia harboring neurons ASH and ASI are located. Red arrowheads indicate fibrotic-like structures observed in mutants. Mutant animals for *mig-6*(*qv33*) display collagen IV fibrotic-like structures and abundant intracellular accumulations (small dotted signals). Loss of function of the TGF-β receptor SMA-6 alone in single mutants *sma-6(wk7)* does not affect collagen IV pattern. In double mutants *sma-6(wk7); mig-6(qv33)*, the loss of *sma-6* enhances the collagen IV phenotypes of *mig-6* mutants. Overexpression of the TGF-β receptor in the wild-type background animals *+; sma-6(+) OE* (using *yxEx615* [*Psma-6::sma-6(+)*]), increases collagen IV build-up phenotypes. **(B)** Percentage of animals displaying collagen IV fibrotic-like structures. **(C)** Quantification of the number of collagen IV fibrotic-like structures observed per animal. **(D)** Quantification of EMB-9::mCherry fluorescence intensity in the head region overlapping with the pharyngeal terminal bulb (indicated with a white rectangle in A, which is the region where the majority of fibrotic-like structures are observed, [67]. **(E)** Quantification of the number of collagen IV fibrotic-like structures observed per animal in control and *sma-6(+) OE*. **(F)** Quantification of EMB-9::mCherry fluorescence intensity in the head region overlapping with the pharyngeal terminal bulb (indicated with a white rectangle in A), in control and *sma-6(+) OE*. **(G-H)** Analysis of young adult animals, where collagen IV crosslinking enzyme peroxidasin-2 is visualized using reporter *qy76* mNG::peroxidasin-2. **(G)** Fluorescence images of the head region (deconvoluted sum projections). Arrows indicate an unidentified ventral structure observed in animals expressing mNG::peroxidasin-2. **(H, I)** Quantification of fluorescence intensity of mNG::peroxidasin-2 in mutant animals (**H**) or in animals overexpressing *sma-6(+)* (**I**). The yellow rectangle in G indicates the region of interest used (selected as it in the collagen IV fibrotic structures area, and it avoids the intense unidentified ventral structure that would mask more subtle differences if included). Error bars are the standard error of the proportion (in B) or of the mean (in C-F, H-I). z-test (B), Wilcoxon Mann-Whitney test (C), ANOVA (D, H), Wilcoxon test (E), t-test (F and I). A.U., arbitrary units. OE, overexpression of wild-type copies. Scale bars, 20 µm.

We next tested whether increased TGF-β pathway signaling by overexpression of the type I receptor SMA-6 would affect collagen IV, as overactivation of TGF-β signaling in vertebrates leads to fibrosis [77–79], and its inhibition attenuates fibrosis [80, 81]. We generated a strain of transgenic animals carrying multiple wild-type copies of *sma-6(+)* (using extrachromosomal array *yxEx615* [*Psma-6::sma-6(+)*] [42]) and the collagen IV reporter. Overexpression of SMA-6 leads to the appearance of collagen IV fibrotic-like structures (**Fig. 2E**) and an increase in overall collagen IV levels (**Fig. 2F**). These findings align with previous reports on the role of the TGF-β pathway in regulating collagen levels in the ECM [82–86].

Similarly, we examined the pattern of a collagen IV crosslinking enzyme, peroxidasin-2 [87, 88], which we reported to be important for collagen IV organization in the context of neuronal maintenance, and which is upregulated in *mig-6* mutants [67] (**Fig. 2G,H**). Using a reporter for peroxidasin-2 (*qy76* mNG::peroxidasin-2 [35]), we found that loss of the TGF-β receptor SMA-6 alone does not alter PXN-2/peroxidasin levels (**Fig 2G,H**). In contrast, the combined loss of the TGF-β receptor and of *mig-6/papilin* in double mutant animals *sma-6; mig-6* leads to increased peroxidasin-2 levels compared to *mig-6* single mutants (**Fig. 2G,H**). We also examined animals overexpressing SMA-6 (using *Psma-6::sma-6(+)*) and found that it did not impact peroxidasin-2 levels (**Fig. 2,I**). Collectively, these results show that the TGF-β pathway modulates collagen IV remodeling in *mig-6* mutants. TGF-β pathway activation promotes collagen accumulation, consistent with its established role in vertebrate fibrosis, while loss of TGF-β signaling exacerbates ECM defects in *mig-6* animals.

### Loss of *mig-6/papilin* function and alteration of the TGF-β pathway affect ECM proteins laminin and fibulin

Collagen IV remodeling is affected in *mig-6* mutants [67], which is further exacerbated by alterations in the TGF-β pathway (**Fig. 2**), known to regulate the expression of several ECM genes in vertebrates [85, 86, 89] and of cuticle collagens and cuticle ECM-associated genes in *C. elegans* [39–41, 90–92]. This prompted us to investigate whether *mig-6/papilin* and the TGF-β pathway might more broadly affect the ECM organization in the context of head neuronal structures. We thus examined laminin, the second major ECM component. Laminins are heterotrimers composed of α, β, and γ chains. *C. elegans* encodes two α subunits (*lam-3* and *epi-1*), one β subunit (*lam-1*), and one γ subunit (*lam-2*) that form two distinct laminin heterotrimers [93, 94]. We constructed mutant strains carrying a reporter for laminin (using *qyIs7* P*lam-1::lam-1::gfp*) [95]. We found that single mutants for *mig-6/papilin* and for *sma-6*/TGF-β receptor have increased laminin levels compared to wild type, whereas double mutants *sma-6; mig-6* exhibited significantly reduced levels, even lower than wild-type animals (**Fig. 3A,B**). Furthermore, in animals overexpressing the SMA-6 receptor (using *Psma-6::sma-6(+)*), laminin levels were also increased (**Fig. 3C**). These results indicate that *mig-6/papilin* is important to regulate laminin levels in the ECM, and that TGF-β pathway dysregulation, by both loss of function and overexpression, alters laminin levels.

**Figure 3.**
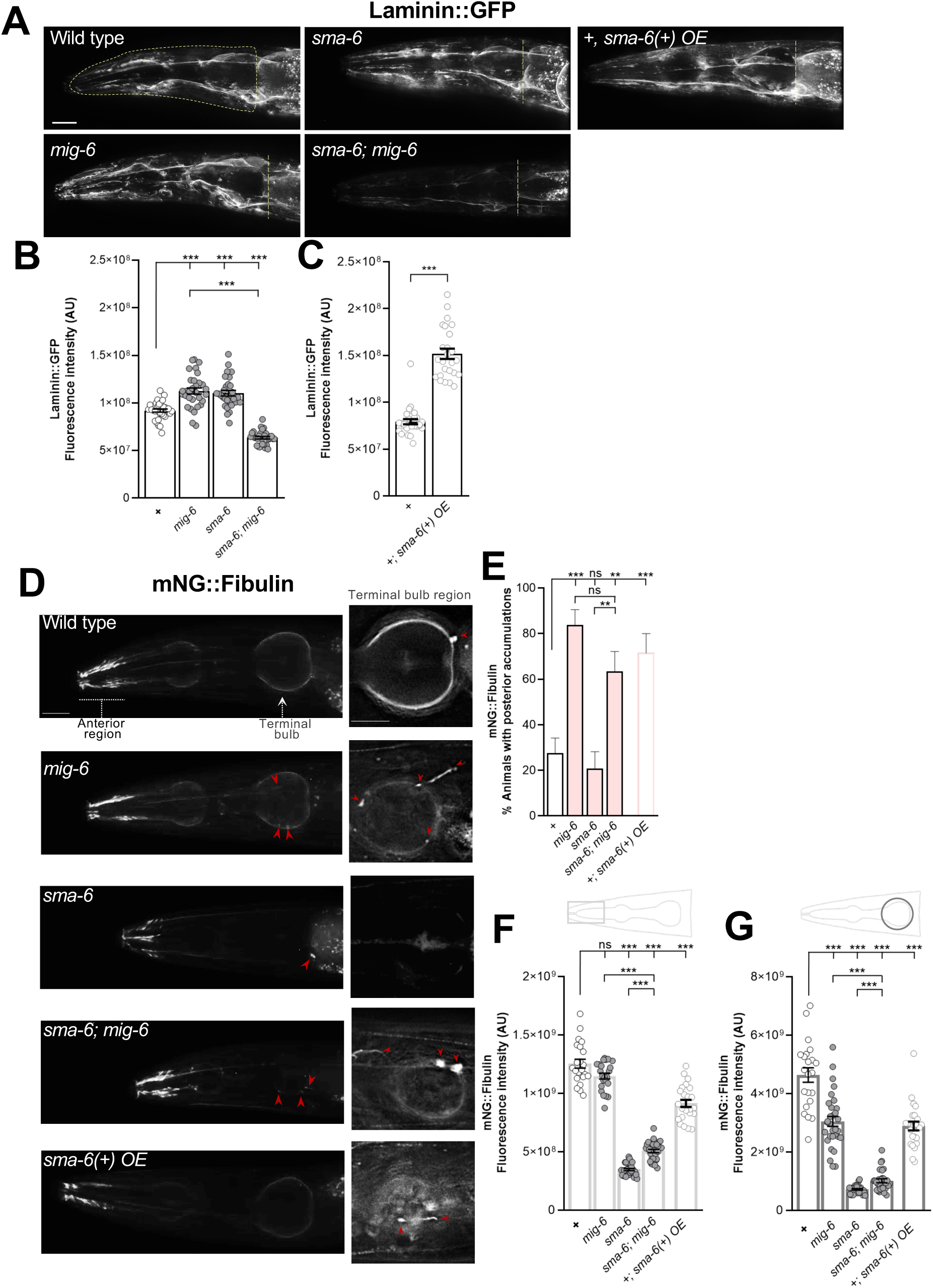
MIG-6/papilin and the TGF-β pathway regulate ECM proteins laminin and fibulin. **(A-C)** Laminin was visualized using reporter *qyIs*7 P*lam-1::lam-1::GFP* in 2-day old adult animals. **(A)** Fluorescence images (deconvoluted sum projections) of the head region showing the ROI used for laminin::GFP signal quantification, outlined in yellow on the wild-type image. For visual clarity, only the posterior limit (yellow line) of the corresponding ROIs is indicated for other genotypes. **(B,C)** Quantification of the fluorescence intensity of laminin::GFP in the head region, as shown in A. **(D-G)** Fibulin was visualized using the mNG::fibulin knock-in reporter *qy62* P*fbl-1::mNG::fbl-1* in 3-day adult animals. **(D)** Left panels are fluorescence images of the entire head (deconvoluted sum projections). Because the mNG::fibulin fluorescence signal is much more intense in the “nose region”, separate sets of animals were imaged using the appropriate acquisition settings to capture the fainter signal in the region of the terminal bulb (right panels; single deconvoluted planes). Examples of fibulin accumulations are indicated by red arrowheads. **(E)** Quantification of the percentage of animals displaying fibulin accumulations in the terminal bulb region. **(F,G)** Quantification of the fluorescence intensity of mNG::fibulin in the nose region (fibulin tracks, pale grey rectangle) and in the terminal bulb region (dark grey circle). Error bars are the standard error of the proportion (E) or of the mean (B, C, F, G). ANOVA (B, F, G), z-tests (E), Wilcoxon test (C). A.U., arbitrary units. Scale bars, 20 µm.

Next, we chose to examine a more dynamic ECM component, fibulin, especially since *mig-6* interacts with *mig-17/ADAMTS* in neuronal maintenance [67], a gene that interacts with the fibulin gene (*fbl-1*) in the context of the developing gonad in *C. elegans* [96–98]. We thus explored if the levels and pattern of fibulin are affected in *mig-6* mutants, as well as in animals with altered TGF-β function, by constructing mutant strains carrying a reporter to examine fibulin (using *qy62* P*fbl-1*::mNG::FBL-1, a knockin reporter that labels all *fbl-1* isoforms [35]. In wild-type animals, fibulin displays strong enrichments in the anterior region of the head, where it lines muscle tracks (**Fig. 3D**; [35]). It is also observed as a fainter signal along the basement membrane of the pharynx and occasionally as spherical accumulations at the posterior ends of the anterior and terminal pharyngeal bulbs (**Fig. 3D-G**). In the examined mutants, fibulin displays complex region-specific changes in both distribution and expression levels. In *mig-6* single mutants, while the anterior head signal is unchanged (**Fig. 3F**), the overall signal at the posterior bulb region is lower (**Fig. 3G**), posterior-bulb accumulations are more frequent (**Fig. 3E**, also on the anterior bulb region, data not shown), and elongated fibrotic-like structures of fibulin occur. In *sma-6* single mutants, fibulin distribution is unaffected, but its levels are strongly decreased (**Fig. 3D-G**). In the double mutant *sma-6; mig-6* animals, the changes in fibulin localization are comparable to those of *mig-6* single mutants (**Fig. 3D,F**), but fibulin levels in the anterior head region and the posterior bulb area are intermediate between those of *mig-6* and *sma-6* single mutants (**Fig. 3F,G**). Finally, overactivation of TGF-β pathway through *sma-6(+)* overexpression (as above) alters fibulin distribution (**Fig. 3D,E**) and decreases its levels (**Fig. 3F,G**). Together, these results show that both *mig-6/papilin* and the TGF-β pathway affect fibulin. Overall, ECM components laminin and fibulin appear to be differently regulated by *mig-6/papilin* and TGF-β signaling.

### Laminin and fibulin are required for neuronal maintenance roles of MIG-6/papilin and TGF-β pathway

We previously showed that collagen IV levels -and its crosslinking-are key for the neuronal stabilizing effect of *mig-6/papilin* mutation [67]. This led us to investigate whether increased laminin levels, observed in *mig-6* mutants, might contribute to *mig-6*-mediated effect on neuronal maintenance. To test this, we depleted laminin by RNAi of *lam-1* or *lam-2* subunits in animals carrying P*sra-6*::DsRed2 to visualize the ASH/ASI neurons. We found that the loss of laminin does not affect ASH/ASI organization (**Fig. 4A**), either in wild-type or in *mig-6* single mutant animals (**Fig. 4A**). However, in *sax-7; mig-6* double mutant animals, depletion of either *lam-1* or *lam-2* leads to the reappearance of *sax-7* neuronal maintenance defects (**Fig. 4A**), indicating that laminin is required for the suppression of *sax-7* neuronal defects implicating *mig-6/papilin* loss of function.

**Figure 4.**
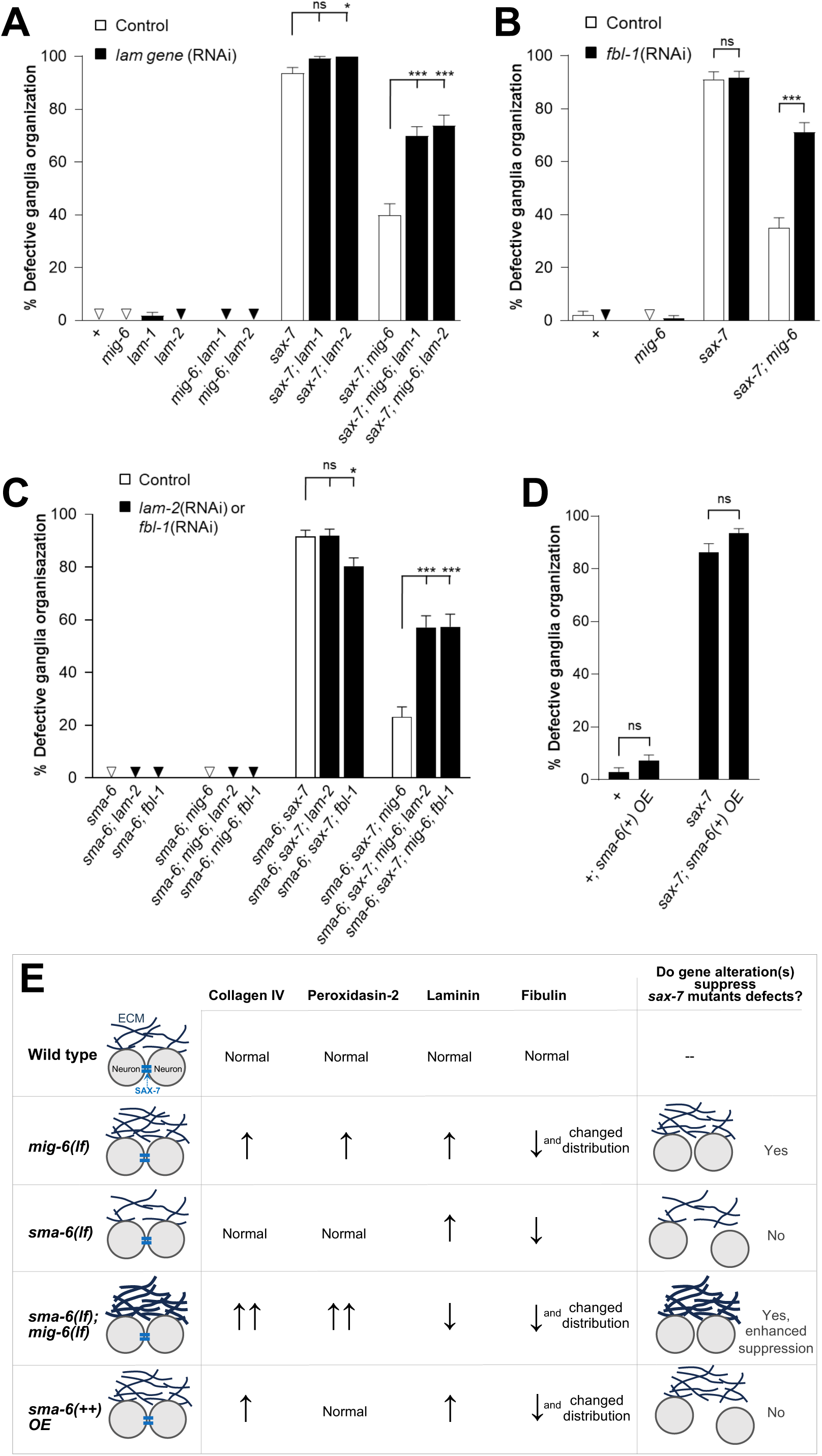
Neuronal stabilization by *mig-6* mutation, and its enhancement through TGF-β signaling inactivation, requires laminin and fibulin. **(A,B)** Quantification of neurons ASH and ASI displacement (using *hdIs26* P*sra-6*::DsRed2, as in Fig. 1) in 2-day old adult animals subjected to control RNAi (empty vector) or to *lam-1(RNAi)* or *lam-2(RNAi)* in **A**, or *fbl-1(RNAi)* in **B**. The suppression of the *sax-7* neuronal maintenance defects by *mig-6(qv33)* mutation is reversed by depletion of laminin or of fibulin. **(C)** Quantification of neurons ASH and ASI displacement in 2-day old adult animals upon depletion of laminin or fibulin by *lam-2(RNAi)* or *fbl-1(RNAi)* in animals lacking TGF-β receptor *sma-6*. The depletion of either laminin or fibulin suppresses the enhancement by loss of TGF-β receptor on the suppression of *sax-7* defects by *mig-6* mutation. **(D)** Quantification of neurons ASH and ASI displacement in 2-day old adult wild-type and transgenic animals overexpressing the TGF-β receptor using *sma-6(+) OE* (as in Figs. 2 and 3). Error bars are the standard error of the proportion. z-tests were performed. **(E)** Summary of the impacts of *mig-6/papilin* loss of function, combined with the loss of the TGF-β receptor, as well as the effect of TGF-β receptor overexpression, on ECM components regulation (increased or decreased levels are indicated by arrows) and on the maintenance of neuronal positioning.

Similarly, we examined the effect of depleting fibulin on neuronal maintenance by knocking down *fbl-1* by RNAi. *fbl-1* depletion does not affect the neuronal phenotype in wild-type animals, nor in single mutants *mig-6* or *sax-7* (**Fig. 4B**). However, in *sax-7; mig-6* double mutants, *fbl-1* knockdown restored neuronal defects (**Fig. 4B**), indicating that fibulin, like laminin, is also involved in the mechanism by which *mig-6* mutation supresses *sax-7* neuronal defects.

Given that TGF-β pathway disruptions affect laminin and fibulin levels and distribution (**Fig. 3**), we next asked if the impact of the TGF-β pathway in *mig-6* mutation-mediated neuronal maintenance (**Fig. 1**) also depends on laminin and fibulin levels. We knocked down *lam-2* or *fbl-1* by RNAi and found that loss of laminin or of fibulin in *sma-6* single mutants, or in *sma-6; mig-6* double mutants, has no effect on neuronal maintenance. *sax-7* defects are modestly reduced by *fbl-1*(RNAi), but not by *lam-1*(RNAi). In contrast, knockdown of *lam-2* or *fbl-1* in triple mutant animals *sma-6; sax-7; mig-6* leads to the reappearance of the neuronal *sax-7* defects (**Fig. 4C**), consistent with the notion that laminin and fibulin may participate in the mechanism by which *sma-6* loss of function enhances the *mig-6-*mutation neuronal stabilizing effect. Finally, we probed if overexpression of *sma-6(+)* alone, which impacted collagen IV, laminin, and fibulin levels/patterns (**Figs. 2** and **3**), could modify the *sax-7* neuronal maintenance defects, and found that it leaves their neuronal disorganization unchanged (**Fig. 4D**). Together, these results suggest that TGF-β pathway disruption alone is not sufficient to induce an ECM state capable of stabilizing neuronal architecture. Rather, the effects of TGF-β inactivation on ECM composition appear to impact neuronal architecture only when combined with the altered ECM state present in *mig-6/papilin* mutants, thereby potentiating ECM remodeling and its impact on neuronal architecture (**Fig. 4E**).

### Disruption of TGF-β signaling pathway alters expression levels of MIG-6S/papilin

Given the genetic interactions between *mig-6/papilin* and the TGF-β pathway documented above, we asked whether the expression levels of papilin protein might be affected by TGF-β signaling disruption. We previously demonstrated that the short isoform of *mig-6/papilin* is responsible for modulating collagen IV remodeling and neuronal maintenance [67]. We thus generated an integrated transgene of the short isoform (P*mig-6*::MIG-6S::mNG) to examine the distribution of MIG-6S/papilin protein in strains with altered TGF-β function carrying this reporter. We examined MIG-6S levels in *dbl-1(nk3)* null mutants [48], as well as upon RNAi-mediated knockdown of the downstream effectors *sma-2* and *sma-3*, and found that loss of the TGF-β ligand or of the TGF-β effectors leads to increased levels of MIG-6S (**Fig. 5A,B**). We further examined MIG-6S levels in animals overexpressing the TGF-β receptor SMA-6 and found that this also increases MIG-6S levels (**Fig. 5C**). Thus, MIG-6S protein levels are affected by both upregulation and downregulation of the TGF-β pathway.

**Figure 5.**
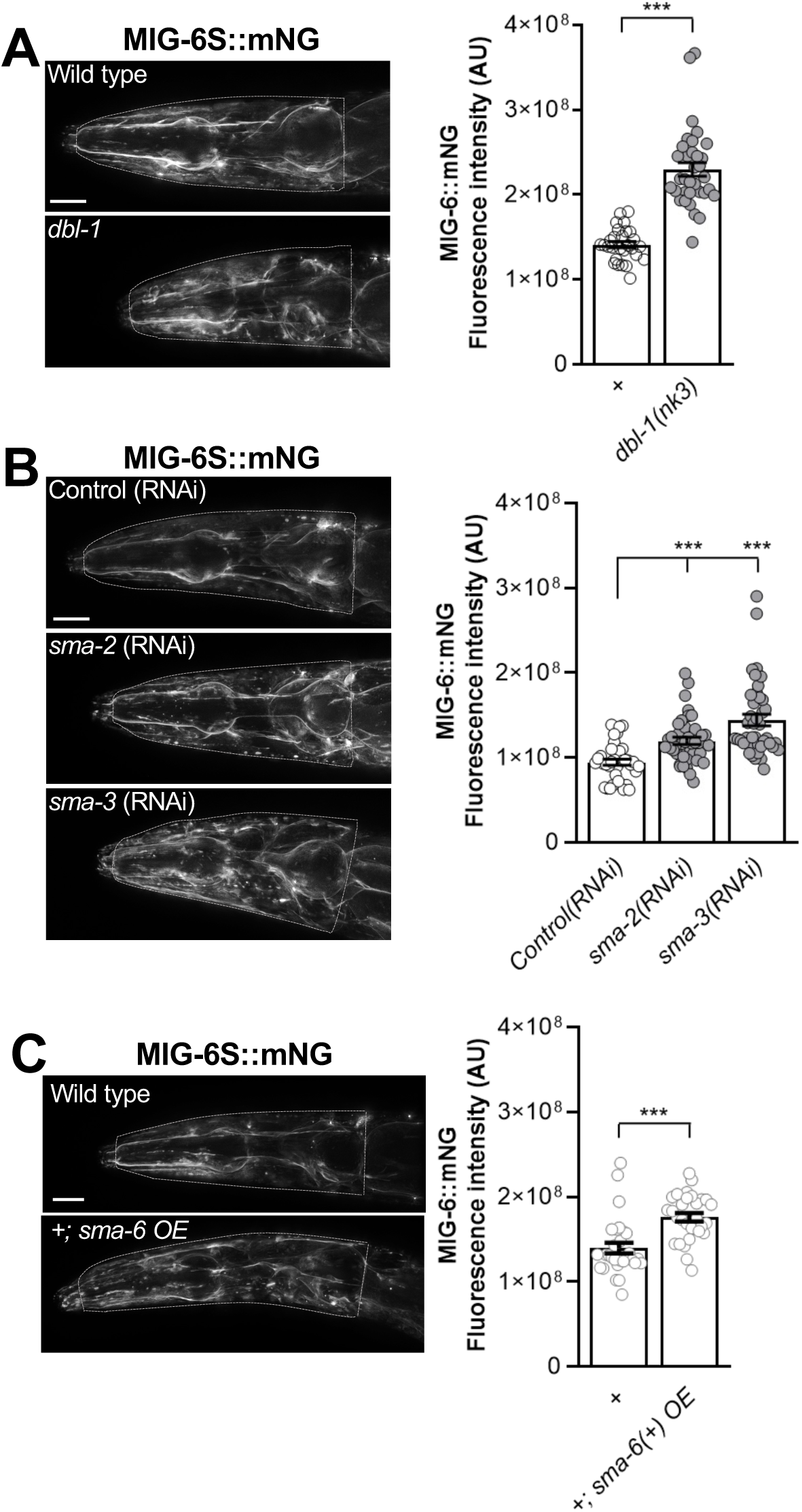
TGF-β pathway loss or overexpression both increase MIG-6S/papilin levels. **(A-C)** Fluorescence images of MIG-6S::mNG using reporter *qvIs9 Pmig-6::mig-6S::mNG* in the head region of 2-day-old adult animals (deconvoluted sum projections). The ROI used for quantification is indicated by the white head contour. **(A)** Quantification of MIG-6S::mNG signal in control and TGF-β ligand *dbl-1(nk3)* mutants, **(B)** in animals subjected to *sma-2*(RNAi), *sma-3*(RNAi), or control RNAi (empty vector), and **(C)** in *sma-6(+) OE* transgenic animals and controls. Error bars are the standard error of the mean. t-test (A, C), ANOVA (B). A.U., arbitrary units. Scale bars, 20 µm.

### Loss of *mig-6/papilin* leads to lowered TGF-β signaling

We next sought out to determine whether TGF-β signaling is altered in *mig-6* mutants, since the inactivation of this pathway potentiates the effect of *mig-6* mutation on neuronal maintenance, suggesting that TGF-β signaling may already be attenuated in *mig-6* mutants. In support of this notion, we noticed that *mig-6* mutants exhibit a reduced body length compared to wild-type animals, as previously noted by [99], which is a characteristic of diminished TGF-β signaling, as this pathway positively regulates body length in *C. elegans* [100, 101]. We measured body length and confirmed that *mig-6* mutants are indeed smaller than the wild type (**Fig. 6A**). To further probe TGF-β pathway activity in *mig-6* mutants we examined the effect of overexpression of the DBL-1 ligand. We used transgene *ctIs40* P*dbl-1::dbl-1(+)* that overactivates TGF-β signaling, with transgenic animals having very long bodies [18, 102]. Whereas *dbl-1(+)* overexpression did increase body length in otherwise wild-type animals, as expected, this increase in body length was abrogated in *mig-6* mutants (**Fig. 6A**). This result indicates that loss of *mig-6* function impairs TGF-β pathway activation by ligand overexpression in this context.

**Figure 6.**
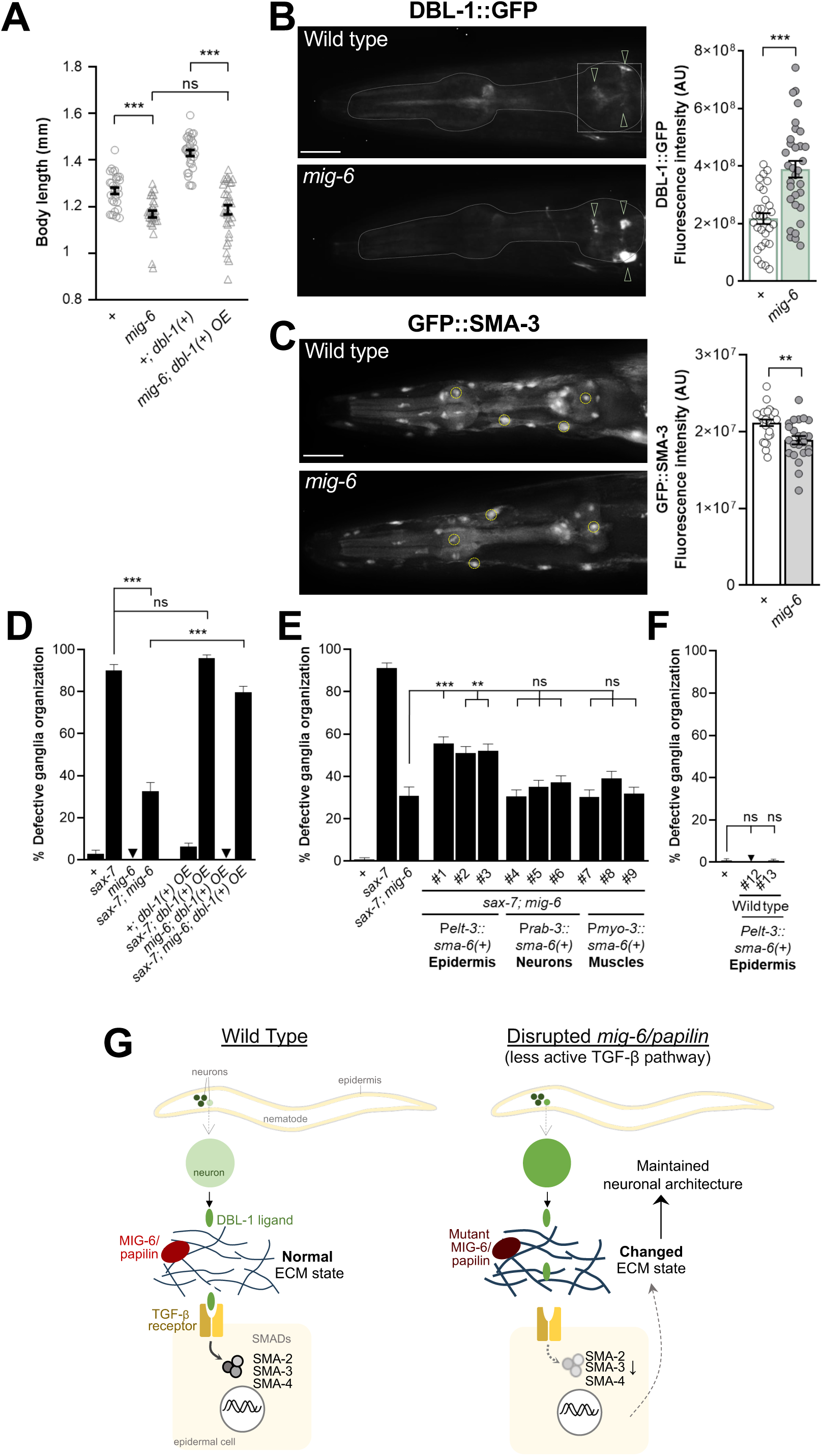
Interplay between *mig-6/papilin* and TGF-β signaling. **(A)** *mig-6* impacts DBL-1/TGF-β-mediated body length regulation. Analysis of body length of 2-day old wild-type and *mig-6* mutant animals, with or without *dbl-1* overexpression [*ctIs40*, *Pdbl-1::dbl-1(+)*].**(B)** Fluorescence images and quantification of DBL-1::GFP of 2-day old control and *mig-6(qv33)* mutant animals. The ROI for DBL-1::GFP signal quantification is indicated (white square). Arrowheads point to signal in DBL-1-expressing neurons. **(C)** Fluorescence images and quantification of GFP::SMA-3 [*qcIs6* (GFP::SMA-3)] of 1-day old adult control and *mig-6(qv33)* mutant animals. ROIs used were four nuclei per animal (small yellow circles); the graph provides the average of signal intensities for four nuclei per animal. **(D-F)** Quantification of neurons ASH and ASI displacement (using *hdIs26* P*sra-6*::DsRed2, as in Fig. 1) in 2-day old adult animals. **(D)** Overexpression of TGF-β ligand *dbl-1(+)* reverses the suppression of *sax-7(qv30)* neuronal defects by loss of *mig-6*. **(E)** Overexpression of TGF-β receptor *sma-6(+)* in the epidermis reverses the suppression of *sax-7(qv30)* neuronal defects by *mig-6(qv33)*. **(F)** Control of overexpression of TGF-β receptor *sma-6(+)* in the epidermis, which leaves neuronal organization unaffected. Error bars are the standard error of the mean (A-C) or of the proportion (D-F). Wilcoxon-Mann-Whitney test (A), t-test (B, C), z-tests (D-F). Scale bars, 20 μm. **(H)** Model for the roles of MIG-6/papilin and of TGF-β pathway in neuronal maintenance. TGF-β ligand DBL-1, produced by neurons, signals through SMA-6/DAF-4 receptors on epidermal cells to regulate ECM gene expression; the resulting levels and organization of the encoded ECM proteins contribute to neuronal maintenance. In *mig-6(qv33)* mutants, the altered state of the ECM (fibrotic, more abundant and more crosslinked) may hinder ligand accessibility to their receptors, thus reducing TGF-β receptor availability at the cell surface, impacting SMAD activation, and ultimately resulting in the inactivation or reduced efficiency of TGF-β signaling.

One possibility for *mig-6* mutants’ smaller body length may be decreased DBL-1 levels. To test this, we measured the levels of the DBL-1 ligand using a well-established reporter transgene for DBL-1 (*texIs100* P*dbl-1::dbl-1::gfp* [23]). We observed that DBL-1, which is known to be expressed in several head neurons [18, 42, 48, 103], shows a stronger signal in these neurons in *mig-6* mutants compared to the wild type (**Fig. 6B**). However, *texIs100* expressing DBL-1::GFP enables to visualize DBL-1 expression inside neurons, while extracellular DBL-1 is not visible, meaning this transgene does not report on the levels of ligand actually reaching the receptor expressed by distant epidermal cells [104]. To better characterize the activation state of the TGF-β pathway in *mig-6* mutants, we therefore measured the levels of the SMA-3 downstream effector using reporter *qcIs6* P*sma-3*::GF::SMA-3 [100]. We found that the levels of GFP::SMA-3 are significantly decreased in *mig-6* mutants compared to wild-type animals (**Fig. 6C**). This indicates that the loss of function of *mig-6/papilin* results in reduced TGF-β signaling activity.

To further confirm that decreased TGF-β pathway activity contributes to *mig-6*-mutation-mediated neuronal stabilization, we tested the effect of overexpressing the DBL-1 ligand (using *ctIs40* P*dbl-1::dbl-1(+)*, [18, 102]. For this, we constructed several strains of single and double mutants of *sax-7(qv30)* and *mig-6(qv33)* carrying this *dbl-1(+)* transgene (and the P*sra-6*::DsRed2 reporter to visualize neurons ASH/ASI). We found that *dbl-1(+)* overexpression does not impact neuronal maintenance in wild-type animals, nor in *mig-6* or *sax-7* single mutants (**Fig. 6D**). However, *dbl-1(+)* overexpression significantly reverses the suppression of the *sax-7(qv30)* neuronal defects by *mig-6(qv33)*, as the penetrance of neuronal defects goes from 30% to nearly 80% in double mutants *sax-7; mig-6* overexpressing *dbl-1(+)* (**Fig. 6D**). This finding indicates that the attenuated activity of TGF-β pathway is part of the neuronal stabilization observed with *mig-6*/papilin loss of function.

In the context of body length regulation, the SMA-6 and DAF-4 receptor subunits function in the epidermis, which is the tissue that secretes the ECM components of the cuticle. To determine in which cell type the TGF-β receptor might function in the context of the maintenance of head neuronal architecture, we drove expression of wild-type copies of *sma-6(+)* by the epidermis, body wall muscles, or neurons, using heterologous promoters, and examined the effect on neuronal architecture. We found that the overexpression of *sma-6(+)* in the epidermis, but not in other cell types, partially reverses the suppression of *sax-7* neuronal defects by *mig-6* mutation. Indeed, double mutant animals *sax-7; mig-6* carrying P*elt-3::sma-6(+)* display approximately 50% neuronal defects compared to the 30% of control double mutant animals *sax-7; mig-6* (**Fig. 6E**). Importantly, the reappearance of neuronal defects is not simply due to *sma-6(+)* overexpression by the epidermis, as controls in the wild-type background does not cause neuronal defects (**Fig. 6F**). Thus, overactivation of the TGF-β pathway counteracts the suppression effect of the *mig-6* mutation specifically, further supporting the notion that TGF-β pathway activity is reduced in *mig-6* mutants.

## DISCUSSION

In our previous work, we demonstrated that the conserved extracellular matrix protein MIG-6/papilin regulates collagen IV remodeling and modulates neuronal architecture. Loss of MIG-6/papilin function leads to fibrotic collagen IV accumulation and altered tissue biomechanics, promoting the stabilization of neuronal structures [67]. In the present study, we explore the underpinnings of the fibrosis-like phenotype observed in *mig-6* mutants, focusing on the TGF-β signaling pathway, a well-established driver of fibrosis in mammals. Our results uncover a mechanism by which MIG-6/papilin affects TGF-β signaling to regulate ECM composition and impact neuronal maintenance, revealing MIG-6/papilin as a novel positive modulator of TGF-β pathway activity.

### The TGF-β pathway functions with *mig-6* to regulate the extracellular matrix

The ECM is continuously remodeled by processes such as synthesis, secretion, modification, and degradation of its components, which is essential for tissue architecture restructuring during development and for maintaining normal organ homeostasis [10, 105]. In addition, the ECM undergoes profound remodeling under pathological situations, including tissue fibrosis and cancer [10, 105]. Here, we extend our analysis of the altered ECM state resulting from loss of *mig-6/papilin* function and reveal interactions with the TGF-β signaling pathway. Loss of *mig-6/papilin* leads to an accumulation of crosslinked collagen IV [35, 67], with increased levels of peroxidasin -the enzyme responsible for crosslinking collagen IV chains [67]. Beyond collagen IV, we also found that other key ECM components are also affected in *mig-6*/*papilin* mutants: laminin accumulates at higher levels within the ECM, while fibulin displays altered distribution patterns and locally reduced levels (**Figs. 2, 3**). Thus, since the loss of function of *mig-6/papilin* leads to a fibrotic-like ECM state, we propose that MIG-6/papilin plays an anti-fibrotic role in ECM remodeling.

Interestingly, disruption of the TGF-β pathway further modifies the *mig-6* mutants’ ECM phenotypes: inactivation of TGF-β in the double mutants *sma-6; mig-6* results in an even greater accumulation of fibrotic collagen IV compared to *mig-6* single mutants (**Fig. 2**). Laminin levels decrease upon TGF-β inactivation (**Fig. 3**), while the distribution of fibulin appears similarly affected as in *mig-6* mutants, depending on regions of fibulin enrichment (**Fig. 3**). Furthermore, overactivation of the TGF-β pathway promotes the accumulation of ECM proteins, including both collagen IV and laminin (**Fig. 2, 3**), consistent with findings in other organisms [106, 107]. Notably, we also observed that MIG-6S/papilin levels are regulated by TGF-β signaling (**Fig. 5**). Altogether, we demonstrate that loss of *mig-6*/papilin leads to a fibrotic-like ECM state analogous to that described in fibrosis, a phenotype that is further exacerbated by the disruption of TGF-β signaling pathway.

Fibrosis, a pathological hallmark of many chronic inflammatory diseases, is characterized by the excessive accumulation of multiple ECM components [108]. In mammalian models, TGF-β ligands are upregulated and activated within fibrotic tissues, playing a key role in driving fibrogenic responses in multiple organs, including the skin, kidney, colon, lungs, liver, and intestines [106]*. mig-6* mutants display an increase in collagen IV, accompanied by a fibrotic phenotype. TGF-β is known to promote fibrosis by enhancing both the synthesis and accumulation of diverse collagens [107], and contributes to their post-translational crosslinking, enhancing their stability [106]. Here, notably, we found that loss of MIG-6 results in accumulations of non-fibrillar collagen IV in fibrotic forms, a phenomenon also documented in various mammalian tissues, including the liver and kidneys, as well as in the basement membrane of *Drosophila* egg chambers [109–114].

Peroxidasin catalyzes sulfilimine bond formation within collagen IV networks, thus consolidating basement membranes [115]. Importantly, peroxidasin has been reported to be highly expressed in fibrotic contexts, including in murine models of kidney fibrosis and non-alcoholic fatty liver disease [116–118]. Conversely, peroxidasin deficiency has been associated with fibrosis regression and improved liver function, likely due to the looser, poorly crosslinked collagen matrix, which facilitates collagen removal [116]. The levels of peroxidasin respond to TGF-β in a context-dependent manner: it is induced by TGF-β in pulmonary and dermal fibroblasts, and in kidney fibrosis [118], but it downregulated by TGF-β1 during TGF-β1-induced epidermal-to-mesenchymal transition [115]. In our study, we observed elevated expression levels of peroxidasin in *mig-6*/*papilin* mutants [67]; **Fig. 2**), indicating that MIG-6/Papilin regulates collagen IV crosslinking in the ECM.

In addition to collagen IV remodeling in *mig-6* mutants, we found that laminin also accumulates in these mutants (**Fig. 3**). Laminin has been reported to be increased in fibrosis in different tissues [119–121]. Interestingly, laminin has been implicated in collagen IV recruitment into the basement membrane [94, 122, 123], raising the possibility that the higher levels of laminin in *mig-6* mutants could contribute to their increased collagen IV accumulation. In contrast, in *sma-6; mig-6* double mutants, laminin levels are decreased may be downregulated in these animals where the TGF-β pathway is both inactivated by the TGF-β receptor mutation.

We also found that *mig-6/papilin* mutants display an altered distribution of fibulin (**Fig. 3**), an ECM protein that interacts genetically with MIG-17/ADAMTS during gonad morphogenesis [97], and that we have found to function in the same pathway as *mig-6* in the context of neuronal maintenance [67]. Fibulin is a dynamic ECM component, as demonstrated in *C. elegans* [35], and has been reported to be elevated in pulmonary tissue fibrosis, where it regulates TGF-β and is proposed to act as a driver of TGF-β-induced fibrosis [124]. Additionally, previous studies in *C. elegans* have shown that *fbl-1* regulates gonad-arm elongation and expansion [125, 126]. Mutations in *fbl-1* bypass the requirement for MIG-17/ADAMTS in directing distal tip cell migration, and fibulin localizes to the gonadal basement membrane in a MIG-17-dependent manner [97]. Furthermore, mutations in *emb-9* (collagen IV) have been shown to suppress *fbl-1* null phenotypes, suggesting that fibulin regulates EMB-9/collagen IV accumulation in the gonadal basement membrane [125]. More recently, fibulin has been identified as a mediator of a basement membrane linkage (B-LINK) supporting the uterus during egg-laying in *C. elegans* [127]. Collectively, these findings imply that fibulin may play a role in stabilizing collagen IV in its fibrotic form in *mig-6* mutants, thereby promoting neuronal maintenance by supporting the ECM structure and integrity.

### MIG-6/papilin regulates ECM remodeling through modulation of the TGF-β pathway

Our findings suggest that TGF-β signaling is attenuated in *mig-6/papilin* mutants, supported by multiple lines of evidence (Figs. 1, 4, 6): (1) *mig-6* mutants exhibit the reduced body length compared to wild-type animals; (2) loss-of-function mutations or RNAi-mediated knockdown of TGF-β pathway components (ligand, receptor, intracellular effectors) enhance the suppression of the *sax-7* neuronal maintenance defects by *mig-6* loss-of-function; (3) overexpression of the TGF-β ligand or receptor—thereby forcibly activating the pathway—counteracts the suppression mediated by *mig-6-*mutation and reinstates *sax-7* neuronal defects; and (4) nuclear levels of TGF-β pathway intracellular cofactor SMA-3 are reduced in *mig-6* mutants relative to wild type, reflecting a decrease in TGF-β pathway activity in *mig-6* mutants.

One possible explanation for the downregulation of TGF-β signaling in *mig-6* mutants is altered diffusion of the DBL-1 ligand [128–130] caused by the ECM’s altered state, with increased matrix components and a more crosslinked collagen IV network [67]. Alternatively, excessive accumulation of ECM components, such as collagen IV, may favor their binding to the TGF-β ligand, thereby competing with the receptor for the ligand availability and reducing pathway activation [131, 132]. Additionally, changes in the biomechanical properties of the ECM in *mig-6* mutants [67] may impair TGF-β signaling, as mechanical cues are known to influence ligand bioavailability and activation [133–135]. For instance, in other systems, the TGF-β ligand is secreted as an inactive form requiring extracellular activation [136–138]. It is conceivable that ECM-degrading enzymes, including MIG-17/ADAMTS that functionally interacts with *mig-6/papilin* [67], may participate in this activation mechanism, thus being impaired in *mig-6* mutants and ultimately leading to lower pathway efficiency.

A key finding from our study is that fibrosis persists in *mig-6* mutants despite attenuation of TGF-β signaling. This observation is unexpected, as TGF-β pathway inhibition typically reduces fibrosis studies in vertebrate models [80, 81, 139]. Our results raise the possibility that the ability of TGF-β signaling to regulate the expression of ECM components may depend on papilin. It will be important to determine whether transcription of ECM components is unchanged despite TGF-β pathway inhibition in *mig-6* mutants, or whether other pathways may drive fibrosis independently of TGF-β, such as MAPK, Wnt/β-catenin, JAK/STAT, or RAS/ERK signaling [140, 141].

It is well documented that TGF-β overexpression promotes fibrosis in both *in vivo* and *in vitro* models [106, 107, 142]. In humans, *TGF-β overexpression induces pronounced fibrotic changes across multiple tissues* [78, 79, 143]. Conversely, loss of one of the downstream effectors, for instance, has been shown to reduce tissue fibrosis in various organs, such as skin [144], kidney [145], lung [146], liver [147], and intestine [148, 149]. The activation of TGF-β signaling through overexpression of TGF-β type I receptor, or of DBL-1 ligand, leads to the reappearance of the *sax-7* mutants’ neuronal defects, specifically in the *mig-6* mutant background. This is consistent with the notion that the TGF-β pathway is less active in *mig-6* mutants, and that the state of TGF-β signaling is critical for the mechanisms by which *mig-6* loss of function suppresses *sax-7* neuronal defects. Based on our findings, we thus propose that MIG-6/papilin may act as a positive modulator of the TGF-β pathway. Further investigation is needed to determine whether the positive effect of MIG-6 on TGF-β signaling is direct or mediated through indirect mechanisms.

MIG-6/Papilin is an ADAMTS-like protein [150, 151]. ADAMTS-like proteins (ADAMTSLs) are structurally related to the ADAMTS family of ECM proteases [152, 153] but lack a catalytic domain [151]. Elevated levels of ADAMTSLs have been reported in fibrotic conditions, including increased ADAMTSL1-5 and papilin in the hearts of mice with fibrosis [150, 154]. Interestingly, ADAMTSL2 has been proposed to function in a negative feedback loop regulating TGF-β signaling, with elevated ADAMTSL2 attenuating TGF-β pathway activity, potentially by acting upstream of active, released TGF-β [150]. This inhibitory role of ADAMTSL2 contrasts with our findings for wild-type MIG-6/papilin function, which promotes TGF-β signaling (as TGF-β signaling is attenuated in loss-of-function *mig-6* mutants). Similar regulatory diversity has been reported for BMP signaling, where both positive and negative regulators have been identified. For example, in *C. elegans*, transmembrane regulators such as DRAG-1 [155], UNC-40/neogenin [156], SMA-10/LRIG [157], CRM-1/CRIM [158], and tetraspanins [159, 160], as well as enzymes SUP-17/ADAM10 and ADT-2/ADAM ([90, 159] act to positively regulate BMP signaling. On the other hand, LON-2/glypican and LON-1 have been described as negative regulators of the pathway [73, 161, 162]. In sum, our study identifies MIG-6/Papilin as a novel extracellular matrix positive regulator of TGF-β signaling.

We also found that neuronal DBL-1::GFP levels are elevated in *mig-6* mutants compared to wild-type animals (**Fig. 6**). As GFP fusion proteins frequently are not reliably detected in the acidic extracellular milieu, it is possible that this reporter only informs on the intracellular levels of DBL-1::GFP. In any case, the observation that DBL-1::GFP levels are elevated in *mig-6* mutants suggests that MIG-6/papilin may be required for normal expression and/or secretion of DBL-1 into the ECM. Alternatively, insufficient DBL-1 delivery to its target may ultimately lead to reduced TGF-β pathway activity, triggering a compensatory feedback mechanism to increase *dbl-1* expression, suggesting a role of MIG-6/papilin in DBL-1 distribution in the ECM. In *C. elegans*, it has been shown that ADT-2/ADAMTS is necessary for normal DBL-1 expression levels to maintain normal cuticle structure and regulate body size [90].

Both the loss of TGF-β signaling and its overactivation -via overexpression of the *sma-6* type I receptor-lead to increased MIG-6S/papilin levels (**Fig. 5**). We propose that under TGF-β pathway overactivation, which induces a fibrotic state, elevated papilin levels may act to compensate and maintain a normal ECM state. Conversely, the increase in MIG-6S/papilin levels observed upon disruption of the TGF-β pathway suggests that TGF-β signaling may normally act to downregulate papilin expression. Interestingly, this is reminiscent of findings that ADAMTSL2 expression is regulated by TGF-β during fibrous tissue differentiation in the sclerotome [163].

### A *mig-6*-dependent role for the TGF-β signaling pathway in neuronal maintenance

We previously showed that collagen IV (both its levels and crosslinking) is required for the maintenance of neuronal architecture mediated by loss of *mig-6*/papilin function [67]. We have now found that laminin and fibulin are also necessary in the mechanism by which the ECM contributes to maintaining neuronal organization in *mig-6* mutants, as knockdown of laminin or fibulin reverses the suppression of *sax-7* neuronal maintenance defects by *mig-6* mutation (**Fig. 3**). We further show that the TGF-β pathway impacts ECM remodeling, which is consistent with its previously described roles in apical ECM regulation in the context of cuticle formation in *C. elegans* [23, 70, 92, 164]. Here, we studied the role of the TGF-β pathway on the ECM that composes the basement membrane ensheathing head neuronal structures, which comprises collagen IV. We found that, in this context, the TGF-β pathway impacts ECM regulation in a *mig-6*-dependent manner: with normal *mig-6/papilin* function, inactivation of the TGF-β pathway only minimally affects the ECM composition in the head region and has no impact on neuronal organization. However, when *mig-6* is mutated, the impact of TGF-β on neuronal maintenance is revealed, with loss of TGF-β signaling pathway components -including of the ligand *dbl-1*, the receptors *sma-6* and *daf-4*, and the downstream effectors-enhancing the suppression of *sax-7* neuronal defects by *mig-6* mutation (**Fig. 1**). Conversely, TGF-β components overexpression has no effect on neuronal organization by itself, but efficiently counteracts the suppression of *sax-7* neuronal defects by *mig-6* (**Fig. 6**).

Thus, we identified a nonautonomous mechanism involving TGF-β signaling in the epidermis that contributes to the maintenance of neuronal organization in *C. elegans*. The neuronally-produced TGF-β ligand DBL-1 is known to signal to peripheral tissues by binding to the heterodimeric receptor complex composed of the type 1 receptor SMA-6 and type 2 receptor DAF-4, and acts via the nucleocytoplasmic SMAD proteins SMA-2, SMA-3, and SMA-4 [18, 40, 48, 165, 166]. Our working model (**Fig. 6G**) proposes that the TGF-β pathway contributes to long-term neuronal maintenance in a *mig-6*-dependent manner: DBL-1 ligand generated from a subset of neurons [42] signals through the SMA-6/DAF-4 receptors in the epidermis to regulate gene expression, including ECM genes impacting neuronal maintenance. As discussed above, in *mig-6* mutants, the TGF-β signaling pathway appears to be less active, which may be attributable to the fibrotic ECM environment in *mig-6* mutants [67], further altering the regulation of ECM remodeling. In sum, our findings reveal a novel mechanism in MIG-6/papilin contributes to ECM remodeling, at least in part through regulation of the TGF-β pathway, and provide molecular insight into how a fibrosis-like state supports the long-term maintenance of neuronal architecture. This work underscores the critical role of the ECM in safeguarding neuronal architecture—an area that remains relatively unexplored.

## Acknowledgements

We thank members of the Bénard laboratory for advice throughout this study and comments on the manuscript; Cathy Savage-Dunn, Tina Gumienny, Meera Sundaran, Kiyoji Nishiwaki, Cassie Blanchette, and Jean-Claude Labbé for stimulating discussions; Grégoire Bonnamour (UQAM) for confocal microscopy expertise; several laboratories for sharing strains and/or plasmids, including D. Sherwood, R. Pocock, Y. Zhang, C. Savage-Dunn, T. Gumienny, A. Fire, and S. Hekimi; Wormbase (www.wormbase.org) provided information about genome sequence and annotations; the *Caenorhabditis* Genetics Center, which is funded by NIH Office of Research Infrastructure Programs (P40 OD010440) for strains; WormAtlas, which is funded by NIH OD010943 to David H. Hall.

## MATERIALS AND METHODS

### *C. elegans* strains and genetics

Strains were cultured at 20°C on nematode growth medium (NGM) agar plates seeded with *E. coli* OP50 bacteria as described [167], unless otherwise specified. N2 is the reference wild-type strain. Mutant alleles used and strains constructed, using standard genetic procedures, are listed in Table S1. Genotypes were confirmed by visible phenotypes when possible, and by genotyping PCR or sequencing, using primers listed in Table S2. All mutant alleles and reporter strains were outcrossed at least three times prior to strain building or analysis.

### Neuroanatomical analyses

The cell bodies of neurons ASH/ASI and their axons, which project into the nerve ring, were visualized using reporter P*sra-6::DsRed2* (*hdIs26*) in 2-day old adult nematodes (selected as late L4 and observed 48 hours later). These animals were mounted on 5% agarose pads, immobilized with 75 mM NaN_3_, and observed under fluorescence microscopes Axio Scope.A1 or Axio Imager.M2 (Zeiss), with a 40x objective. Images were acquired using an AxioCam camera (Zeiss) and processed using ZEN (Zeiss). In wild type animals, the ASHL/R and the ASIL/R soma are located posterior to the nerve ring. An animal was counted as mutant when at least one of the ASH/ASI soma, along the antero-posterior axis of the animal, was not posterior to the nerve ring (but either anterior to or flanking the nerve ring).

### Analysis of Collagen IV fibrotic structures

2-day old adults (selected as late L4 and observed 48 hours later) were mounted on 5% agarose pads and immobilized with 75 mM NaN_3_ and observed under fluorescence microscope Axio Imager.M2 (Zeiss), with a Plan Apo 40x/0.95 NA objective. Images were acquired using an AxioCam camera (Zeiss) and processed using ZEN (Zeiss). Fibrotic collagen IV are enrichments of EMB-9::mCherry signal present in the posterior head region of *mig-6(qv33)* mutant animals [67]. These structures are virtually never observed in the wild type and are different from the wild-type pattern of collagen IV signal present along the body wall muscles basal lamina. To assess the presence of collagen IV fibrotic-like structures in each animal, z-stacks were captured, and all the z-planes images were examined. Elongated structures of collagen IV accumulation that were present in the posterior part of the head, and that are distinct from the faint basement membrane signal along body wall muscles, were considered to be collagen IV fibrotic structures [67]. The number of fibrotic structures per genotype was also counted.

### Microscopy and fluorescence intensity quantification

Animals were observed at precise ages: as “young adults” (animals that just molted from the L4 stage, <3 hours after the L4 molt), “2-day old adults” (48 hours after the young adult stage), or “3-day old adults” (72 hours after the young adult stage). They were mounted on 5% agarose pads with 75 mM NaN_3_, and images were acquired as a z-stack using a Plan Apo 40x/0.95 NA objective on a Zeiss Axio Imager.M2 (equipped with an AxioCam camera and ZEN software). *AutoQuant X deconvolution* software was used to remove blur and enhance contrast and resolution. A Nikon A1 laser scanning confocal microscope (equipped with an EMCCD camera and NIS elements software) was also used for image acquisition and analysis, using either a Plan Apo λ 40x/0.95 NA or Plan Apo λ 60x/1.4 NA oil immersion objective.

The fluorescence intensity of EMB-9::mCherry, mNG::PXN-2, laminin::GFP, mNG::fibulin, MIG-6S::mNG, DBL-1::GFP, and GFP::SMA-3 signals were quantified using ImageJ software as integrated densities (IntDen) of sum intensity z-projections. For each fluorescence reporter, the appropriate imaging channel was used with all image acquisition parameters being fixed across all genotypes examined for a given reporter, including exposure time, excitation intensity, and gain, taking great care to avoid saturation. The region of interest (ROI) containing mCherry, GFP, or mNG fluorescence signals on the images was outlined manually for whole head, or by using a desired ROI (as shown on each figure). The background signal was determined by choosing four ROIs from regions adjacent to the head but located outside of it; the mean fluorescence of the four background ROIs was subtracted from signal intensity. The following formula was used to correct for background fluorescence: Corrected IntDen = IntDen of experimental ROI - (mean of background ROI * area of experimental ROI).

### Transgenes and rescue assay experiments

#### Pelt-3::sma-6 (pRG62)

This plasmid was generated by Padgett lab [21] and provided by Roger Pocock’s lab (Monash University).

#### Prab-3::sma-6 (pCB503)

The *sma-6* cDNA sequence [2649 bp] was amplified from pRG62 [*Pelt-3::sma-6::GFP*] using primers oCB2370 (CATGATACCGGTTAAGATTGATTGGTGGCTG) and oCB2388 (CATGATGGATCCGTTGAAAAAATGAACATCACC) to add AgeI and BamHI sites, and digested. The *sax-7S* cDNA was removed from plasmid pCB428 (P*rab-3::sax-7S*, [56] by digesting with AgeI and BamHI; the released vector backbone was then ligated with the *sma-6* cDNA.

#### Pmyo-3::sma-6 (pCB504)

The *sma-6* cDNA sequence [2649 bp] was amplified from pRG62 [*Pelt-3::sma-6::GFP*] using primers oCB2389 (CATGATTCTAGAAGTTGAAAAAATGAACATCACC) and oCB2390 (TACGATCTCGAGTTAAGATTGATTGGTGGCTG) to add XbaI and Xhol sites, and digested. The *sdn-1* cDNA was removed from plasmid pCB423 (P*myo-3::sdn-1*, [168] by digesting with XbaI and XhoI; the released vector backbone was ligated with the *sma-6* cDNA. All inserts of finalized clones were verified by sequencing.

Transgenic animals were generated by standard microinjection techniques [169]. For *sma-6* rescue assays by expression of *sma-6(+)* from different tissues, plasmid pRG62 [*Pelt-3::sma-6*] was injected at a concentration of 10 ng/μL with *lgc-11::gfp* as coinjection marker; pCB503 [*Prab-3::sma-6*] was injected at 25 ng/μL with *lgc-11::gfp* as a coinjection marker; and pCB504 [*Pmyo-3::sma-6*] was injected at 1 ng/μL with *lgc-11::gfp* as coinjection marker. All coinjection markers were injected at 50 ng/uL. pBSK(+) was added to each mix to reach a final DNA concentration of 200 ng/µL. To generate transgenic animals in the *sax-7(qv30); mig-6(qv33)* background, DNA mixes were injected into a strain where the *mig-6(qv33)* mutation is balanced [*sax-7(qv30); mig-6(qv33)/dpy-11(e224) oyIs14*]. Wild-type background *hdIs26* was injected to generate control transgenic strains. All transgenic strains are listed in Table S1.

#### RNA interference assays

RNAi experiments were performed by the feeding method using RNAi bacterial clones from the Ahringer RNAi library [170, 171], whose identities were verified by sequencing. Single colonies for the L4440 empty vector negative control, or for clones targeting *dbl-1, daf-4, sma-2, sma-3, sma-4, lam-1, lam-2,* or *fbl-1* were obtained on LB plates with ampicillin (75 μg/mL) and tetracycline (12.5 μg/mL). They were then grown in LB medium with ampicillin (75 μg/mL) for 16 hours at 37°C, to which 1 mM IPTG was added and incubated for an additional hour to induce dsRNA expression (2 mM IPTG was used in the case of *fbl-1* RNAi experiments). Concentrated cultures were seeded onto NGM plates containing 75 μg ampicillin and 1 mM IPTG, and then left to dry at room temperature for overnight induction. For all RNAi clones (*dbl-1, daf-4, sma-2, sma-3, sma-4, lam-1, lam-2, fbl-1*, or empty vector control), synchronized L1 larvae [172] were distributed onto RNAi plates (control and RNAi plates), incubated at 20°C, and examined as at 2-day old adults (48 hours post L4).

#### Quantification of body length

2-day old adult nematodes (selected as late L4 and observed 48 hours later) were mounted on 5% agarose pads, immobilized with 75 mM NaN_3_. Body length was measured, from the mouth opening to the anus on DIC images by using the measurement tool in Axio Imager.M2 (Zeiss).

#### Statistical Analysis

All statistical analyses were performed using R (version 4.1.2) using the R Stats package (’stats’ version 4.4.2). Data are presented as mean ± standard error of proportion, or as mean ± standard error of the mean. As indicated in each Figure Legend, statistical tests were performed using z-test, unpaired two-tailed Student’s t-test, or one-way ANOVA when applicable (for parametric datasets). For non-parametric datasets, the Wilcoxon-Mann-Whitney test was used. Appropriate *post-hoc* tests were performed for multiple comparisons: Bonferroni correction was applied after z-tests and Wilcoxon-Mann-Whitney tests, while Tukey HSD correction was applied following ANOVA. All sample sizes and raw data are available in **Supplementary Information**.

---

**Table S1.**
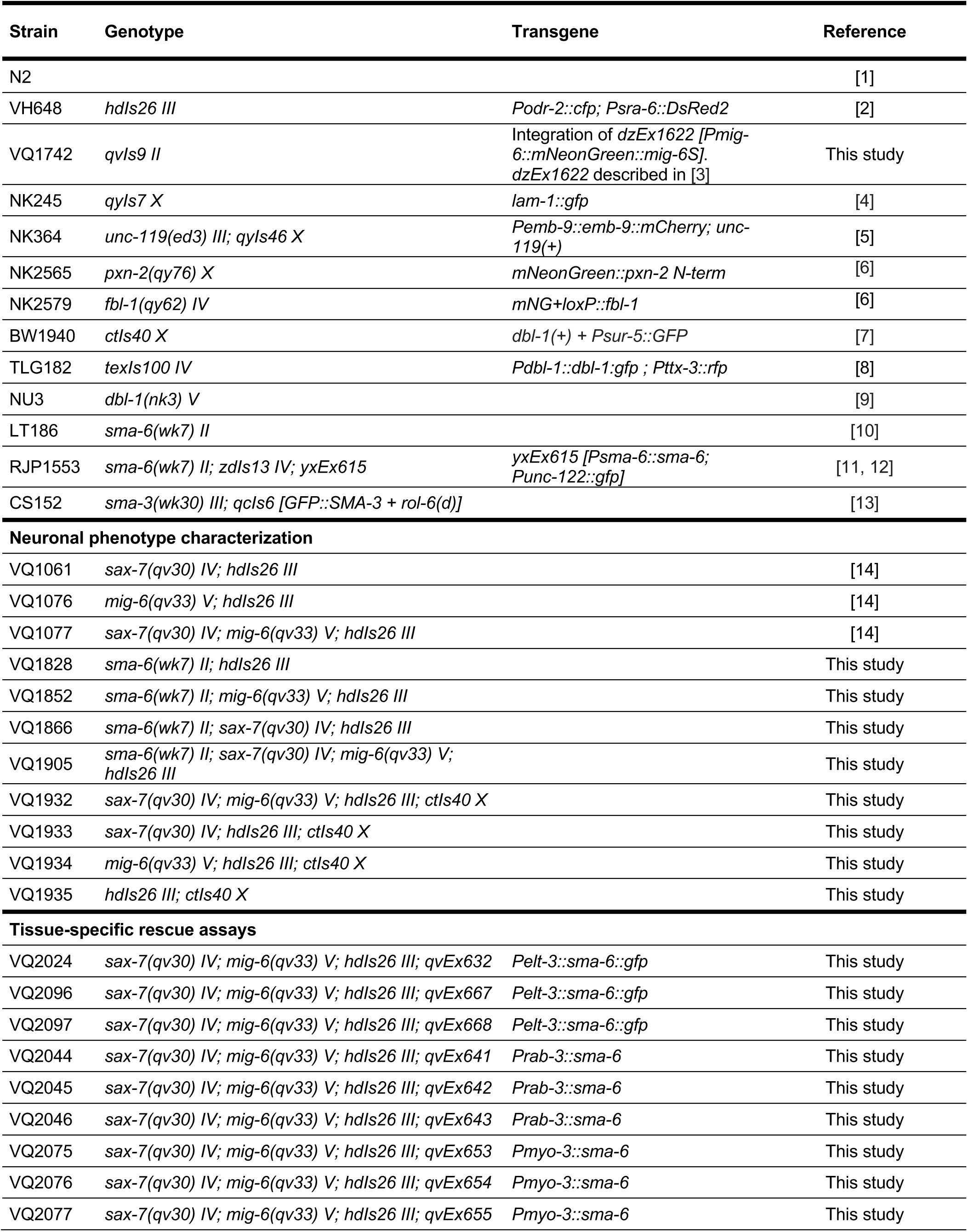
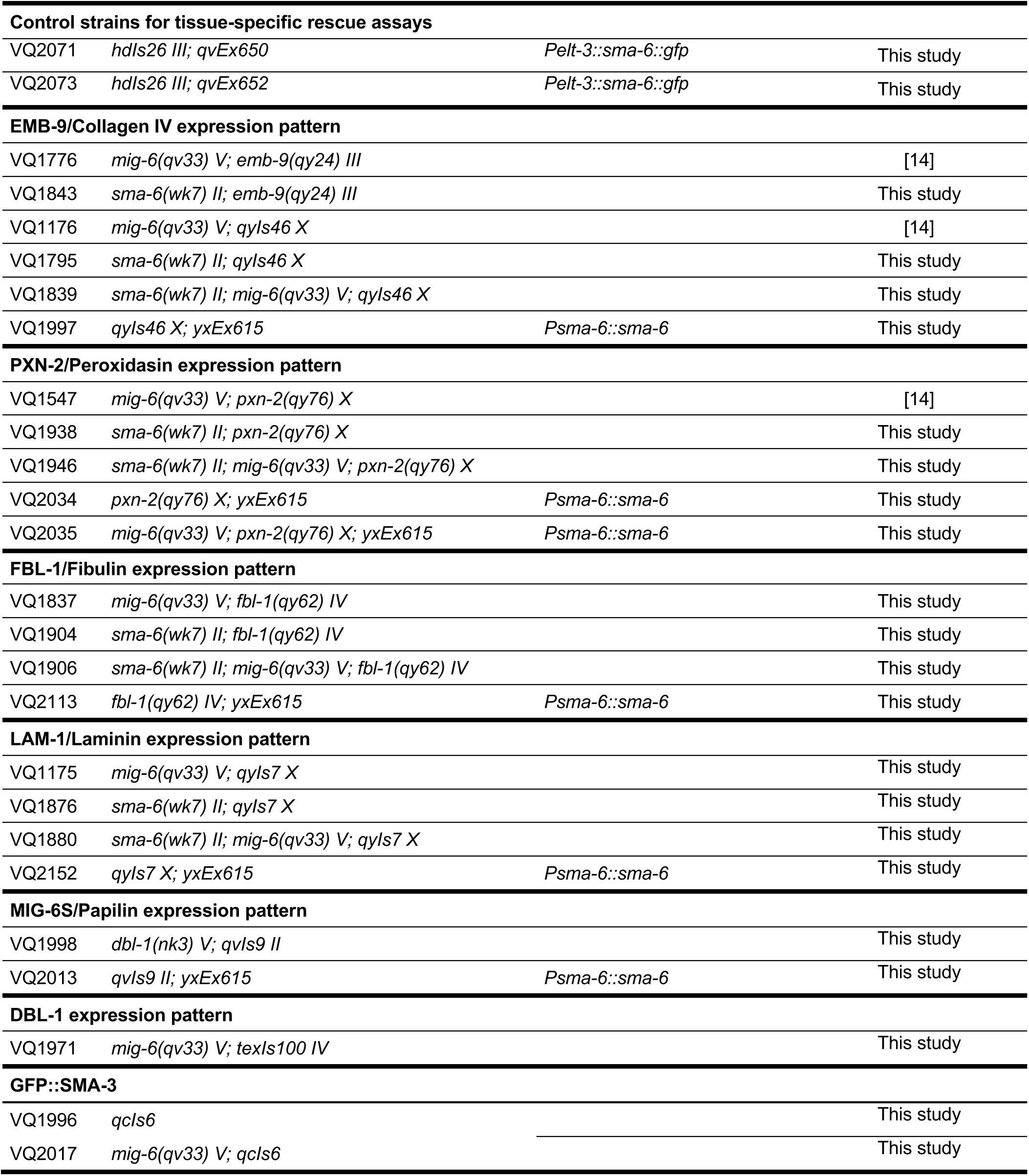
List of strains used.

**Table S2.**
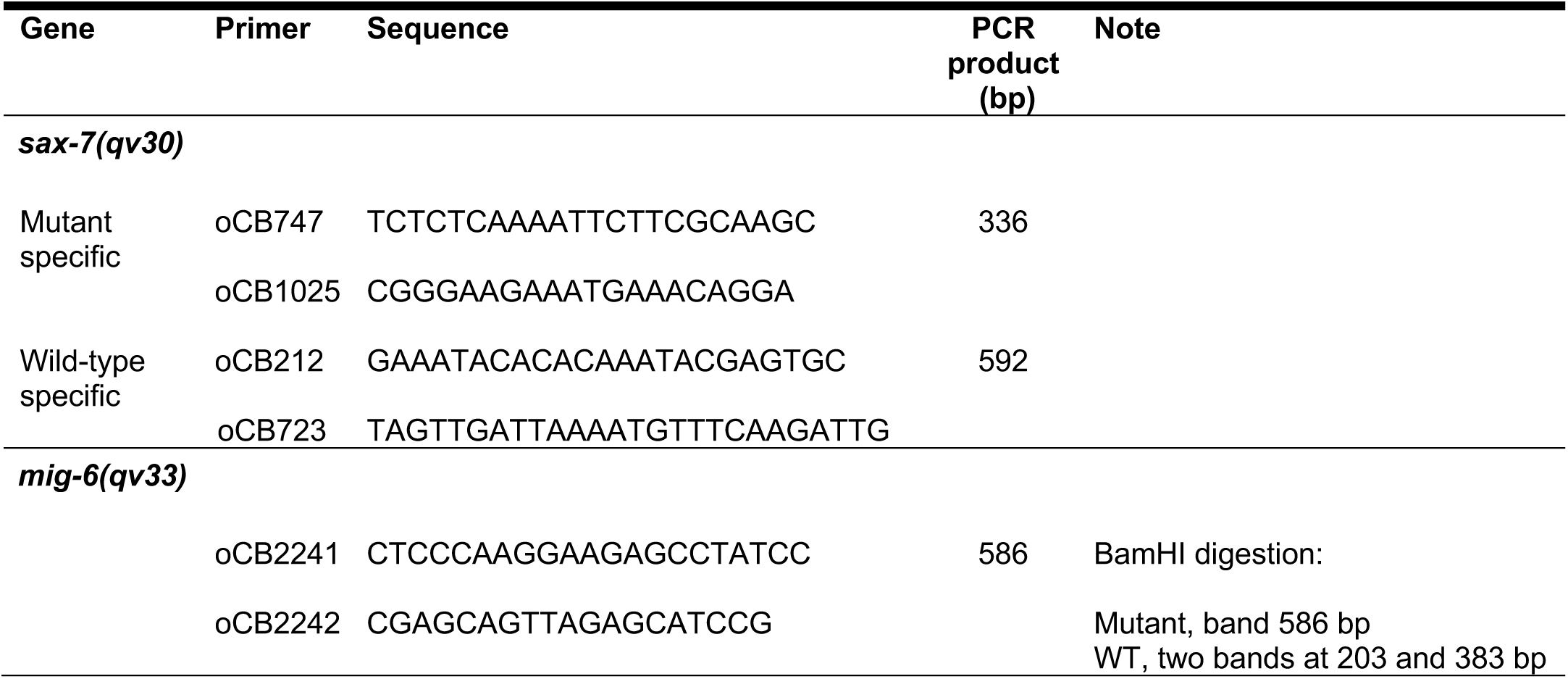
List of primers used.

